# Development and cell cycle dynamics of the root apical meristem in the fern *Ceratopteris richardii*

**DOI:** 10.1101/2020.08.27.271049

**Authors:** Alejandro Aragón-Raygoza, Alejandra Vasco, Ikram Blilou, Luis Herrera-Estrella, Alfredo Cruz-Ramírez

## Abstract

Ferns are a representative clade in plant evolution although underestimated in the genomic era. *Ceratopteris richardii* is an emergent model for developmental processes in ferns, yet a complete scheme of the different growth stages is necessary. Here, we present a developmental analysis, at the tissue and cellular levels, of the first shoot-borne root of Ceratopteris. We followed early stages and emergence of the root meristem in sporelings. While assessing root growth, the first shoot-borne root ceases its elongation between the emergence of the fifth and sixth roots, suggesting Ceratopteris roots follow a determinate developmental program. We report cell division frequencies in the stem cell niche after detecting labeled nuclei in the root apical cell (RAC) and derivatives after 8 hours of exposure. These results demonstrate the RAC has a continuous mitotic activity during root development. Detection of cell cycle activity in the RAC at early times suggests this cell acts as a non-quiescent organizing center. Overall, our results provide a framework to study root function and development in ferns and to better understand the evolutionary history of this organ.

**Summary Statement:** In the Ceratopteris root, the apical cell and its derivatives have a high division frequency, suggesting the apical cell acts as a non-quiescent organizing center in the stem cell niche.

## INTRODUCTION

Exploring the diversity of plant lineages using evo-devo approaches provides insights of how different organs and certain innovations that integrate the sporophyte plant body emerged during the evolution of embryophytes (**Figure 1A**; Delaux et al., 2019). Delving into root evolution, several studies suggest that convergent evolutionary events took place in both extant tracheophyte lineages: lycophytes and euphyllophytes (ferns and seed plants; **Figure 1A, blue circles**). The evolution of roots in lycophytes occurred in a stepwise manner, where gradual stages were found in the fossil record (Fujinami et al., 2020; Hetherington and Dolan, 2018; Kenrick and Strullu-Derrien, 2014; Raven and Edwards, 2001;). Root evolution in lycophytes is considered the first appearance of this organ in vascular plants, followed by a second evolutionary event which likely occurred in the ancestor of euphyllophytes (**Figure 1A**; Liu and Xu, 2018).

**Figure 1.**
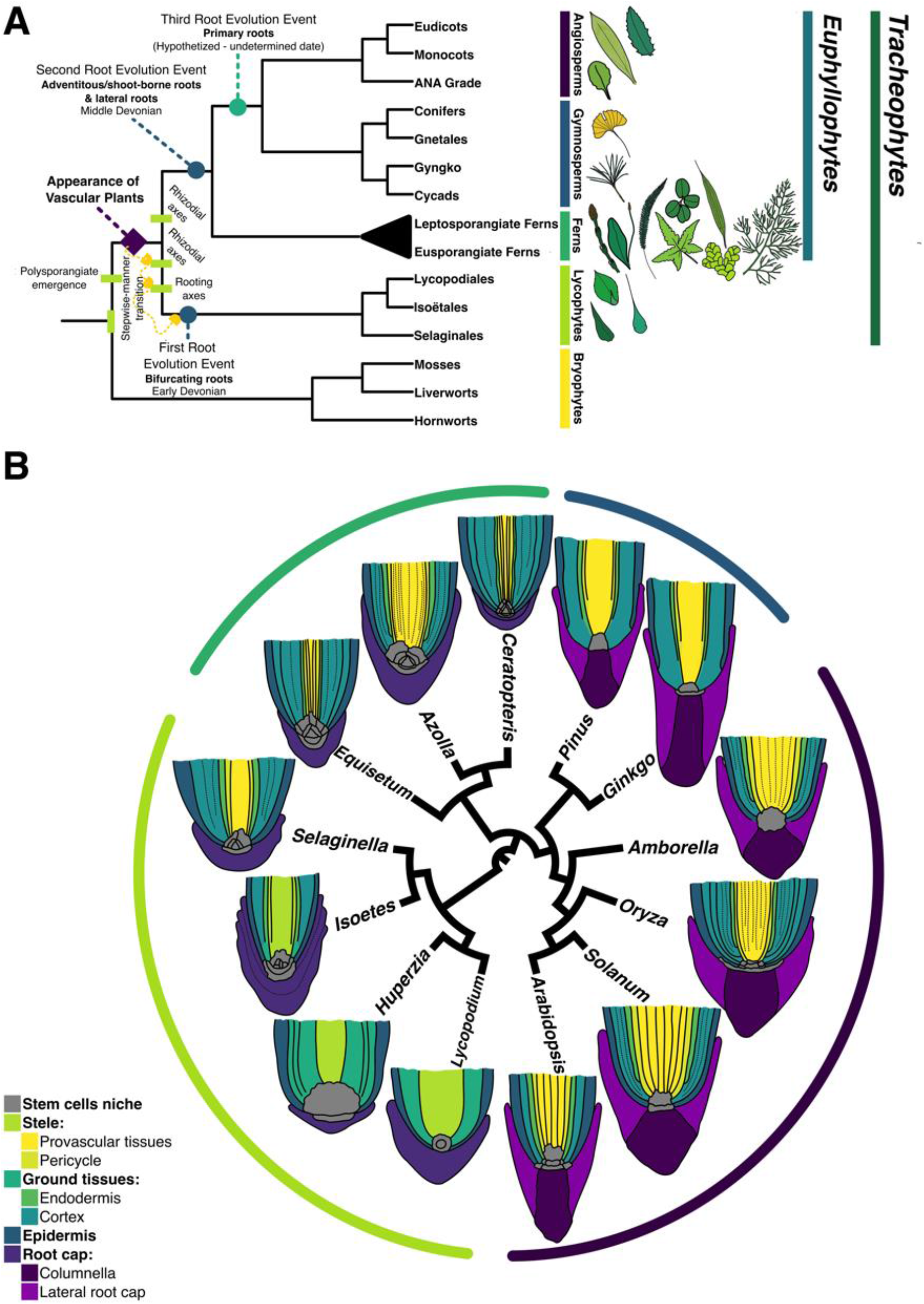
Evolution among major vascular plant lineages and its relation with root meristem organization. **(A)** Hypothesis of root evolution in vascular plants. Both major extant vascular plant lineages, lycophytes and euphyllophytes, have been hypothesized to independently evolve roots. Based on Hetherington and Dolan, 2017, 2018a, 2018b, 2019; Kenrick, 2013; Kenrick and Strullu Derrien, 2014; Liu and Xu, 2018. **(B)** Schematic representation of cell layer organization and stem cell niches in roots from different extant vascular plants (Table S1). Meristem diversity has been extensively assessed across flowering plants, but not in other lineages. *Color lines covering the roots, correspond to color patterns represented in* **A**.

Ferns are the sister group of seed plants and therefore a key lineage to understand the evolution and development of the plant body in tracheophytes (Vasco et al., 2016). *Ceratopteris richardii* (Ceratopteris) is a subtropical fern in the Pteridaceae family, which has been considered “the *Arabidopsis* of the fern world”. This fern presents certain advantages: easiness to culture in laboratory; short life cycle; genetic transformation techniques; and a draft genome sequence (Bui et al., 2014; Hickok et al., 1995; Marchant, 2019; Marchant et al., 2019; Plackett et al., 2014, 2015). Ceratopteris is becoming a more approachable plant model to study developmental biology in ferns. Therefore, an accurate and complete characterization of its ontogenesis is necessary to pursue further research with this organism. Recently, a description summarizing several developmental stages in Ceratopteris was generated (Conway and Di Stilio, 2020). However, a detailed description of the early sporophyte development, immediately after gametophyte fertilization, and root emergence has not been accomplished yet.

Ceratopteris establishes an homorhizic root system where no embryonic root is developed but the continuous emergence of shoot-borne roots. Ceratopteris not only generates stem-borne roots (SBRs), but also leaf-borne roots (Hou and Hill, 2002). SBRs develop from a direct derivative (merophyte) of the shoot apical cell. This derivative will be a progenitor cell for a leaf and a SBR. Leaf-borne roots emerge at the base of each leaf, and more than one root can develop (Hou and Hill, 2002). Some aspects of Ceratopteris root system have been previously characterized, moreover there are several elements from the emergence and development of this organ that need to be determined in order to establish Ceratopteris SBR as a model for developmental biology and evo-devo approaches.

As an organ with active growth, a root bears an apical meristem with stem cells that are mitotically active, to self-renew, and generate specific daughters for diverse cell layers (Dolan et al., 1993; Heimsch and Seago, 2007). The development and arrangement of apical meristems in ferns is one of the most peculiar traits in this lineage. Several fern species display a root apical meristem with a prominent tetrahedral apical cell in the promeristem or stem cell niche (**Fig. 1B; Table S1**; Schneider, 2012). This single apical cell generates all the different cell layers that conform the root body (Hou and Hill, 2002; Hou and Hill, 2004). This is a major difference with the Arabidopsis root stem cell niche (RSCN), where specific initials generate each of the diverse cell layers (Scheres et al., 1995). A similar RSCN to that of Ceratopteris is present in several species of the lycophyte order Selaginellales (**Figure 1B**). This organization is not present in other lycophytes, since other types of RAM arrangement have been described in the orders Isoëtales and Lycopodiales (**Figure 1B**; Fujinami et al., 2017; Fujinami et al., 2020).

In the present work, we performed a developmental analysis of the first stem-borne root of Ceratopteris, at organ, tissue, and cellular levels. We followed the early stages of the embryo and the root meristem establishment after gametophyte fertilization. We detected the diverse set of cell layers that compose Ceratopteris root body by tracking layer-specific traits, such as Casparian strip in endodermis and lignin accumulation in xylem. We analyzed the cell division frequency in the root apical cell and its derivatives at the RSCN after exposure to EdU at different times, and observed a high frequency of labeled S-phase cells after 8 hours of exposure, suggesting that the entire Ceratopteris RAM has an active cell cycle until it enters a determinate program leading to root growth cessation. Our results point out that the root apical cell in Ceratopteris regularly divides which support the hypothesis of this cell high mitotic rate, a disputed fact in ferns development, and propose the absence of a quiescent center in Ceratopteris RSCN.

## RESULTS

### Establishment of the first root during embryonic development in Ceratopteris sporophyte

We explored the early stages of Ceratopteris sporophyte to understand the embryo development and to track the emergence of the first root. While this work was in progress, the different stages of Ceratopteris development were described by Conway and Di Stilio (2020). We not only covered similar stages to those reported, but also expanded the developmental phases of the early sporophytes during zygote formation and embryo development (**Figs. S1A**).

Our results showed that both gametophytes reach their sexual maturity at 15 days post sowing (dps) at 25 °C (**Figs. S1A,B**). Male gametophytes are filled with antheridia and circularized sperms are observed inside them (**Figs. 2A; S1A**). Hermaphrodite gametophytes develop both antheridia and archegonia. Archegonia develop below the notch meristem, and an egg cell develops within each archegonium (**Figs. 2B,C; S1B**). Mobile sperms are able to reach the archegonium tip to fertilize the single egg cell (**Figs. 2D; S1C**). Following fertilization, the zygote forms and then embryo development proceeds (**Figs. 2E-M**).

**Figure 2.**
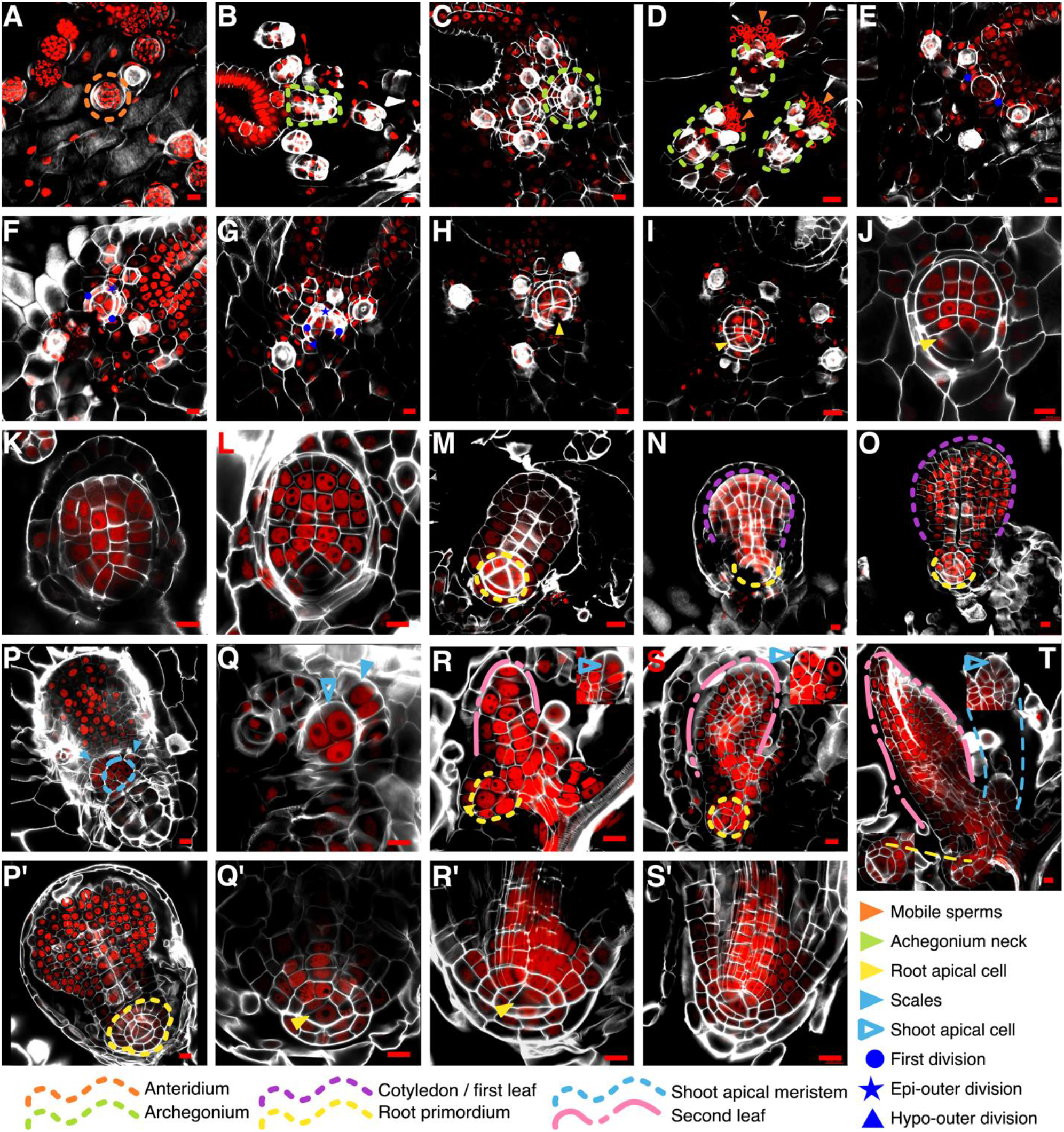
CLSM images of gametophyte-to-sporophyte transition and embryo development in *Ceratopteris richardii*. (**A**) Antheridia in a male gametophyte at 15dps. (**B & C**) Archegonia in a hemaphrodite gametophyte at 15 dps. (**D**) Archegonial necks are reached by mobile sperms. (**E to G**) Early developmental stages of the embryo: two-celled embryo (E); epibasal division in an octant embryo (F); hypobasal division in an octant embryo (G). (**H to O**) The RAC is specified early in embryo development and a root primordium begins to form. Also, the first leaf grows until breaking the calyptra. (**P to T; P’ to S’**). Other components are also developing: shoot apical meristem (P,Q); first root primordium with a root cap (P’,Q’). Sporophytes continue to develop and new organs emerge (R to T). Also, the first SBR begins to grow and the whole root body develops; all cell layers recognizable (R’ & S’). *Scale bars: 20 μm. White – Calcofluor White; Red – Propidium Iodide.*

Between 1 to 3 days after fertilization (daf; **Figs. S1D-F**), the embryo is detected between its two-celled and octant stages (**Figs. 2E-G**). The first division (**Fig. 2E, blue dots**) occurs perpendicular to the anterior-posterior axis of the gametophytes (**Fig. S2**), forming the epibasal (anterior) and hypobasal (posterior) cells. The subsequent division generates an embryo in a quadrant stage, but this division occurs in a vertically median plane and we were not able to observe it (Johnson and Renzaglia, 2008; Wardlaw, 1955). We detected divisions in the outer cells of the quadrant embryo: first, a symmetric division occurs at the epibasal outer quadrant (**Fig. 2F, blue star**), and later an asymmetric division takes place at the hypobasal outer quadrant cell (**Fig. 2G, blue triangle**). The hypobasal outer octants displayed an observable asymmetry, from these octants the first root would emerge (**Fig. 2G, blue triangle**).

At 3 daf, the embryo is recognizable as a swelling lump in the fertilized archegonium (**Figs. 2F; 2G-J, purple arrowhead**), but keeps developing inside the gametophyte, covered by a calyptra (Conway and Di Stilio, 2020). Parallel in time, a prominent tetrahedral-shaped cell can be distinguished at the distal part of the embryo (**Figs. 2H-J; yellow arrowhead**). Because of its position and consistent presence in latter stages (**Figs. 2K-O**), it is likely that this cell is the first root apical cell (RAC). From 4 to 7 daf, the cotyledon (or first leaf) grows and expands, acquiring a rounded shape (**Figs. 2N,O; purple line; FigS1G-J**) and a complete root primordium develops (**Figs. 2N,O; yellow line**), but the growth of each organ occurs at a different rate.

The calyptra breaks around 8-9 daf because of the growing cotyledon (**Figs. S1K,N)**. The cotyledon reaches a wide spatula shape (**Figs. S1N,O, purple line & blue arrowhead**). Around 15 daf, the first SBR begins to grow and a prominent root tip can be clearly observed (**Figs. S1N, yellow line**). All root cell layers can be distinguished after 20 daf (**Figs. S1O, yellow line**). At the same time, the second leaf (**Figs. 2R-T, pink line; S1O**) and a new root primordium appear, and again the growth rate of the root seems to be delayed compared with that of the leaf (**Figs. 2R-T**, **yellow line**). While the first SBR keeps developing, the different layers forming the root body, including the root cap, are detected in longitudinal planes (**Figs. 2P’-S’, blue-filled arrowheads; Fig. S3**).

The cotyledon and the embryonic root developed at a dorsal plane in relation to the gametophyte, the shoot apical meristem (SAM) develops at the ventral plane (**Figs. 2P, blue line**). The SAM is surrounded by scales (**Figs. 2P, blue-filled arrowheads**) and the shoot apical cell (SAC) is distinguishable, due to its inverted tetrahedral shape with a rounded distal face (**Figs. 2P; 2R-T, insets**). Based on observation of multiple events, we summarized Ceratopteris development using a graphic representation from the spore to young sporophytes (or sporelings), with a main focus in the first root specification and development (**Fig. 3**).

**Figure 3.**
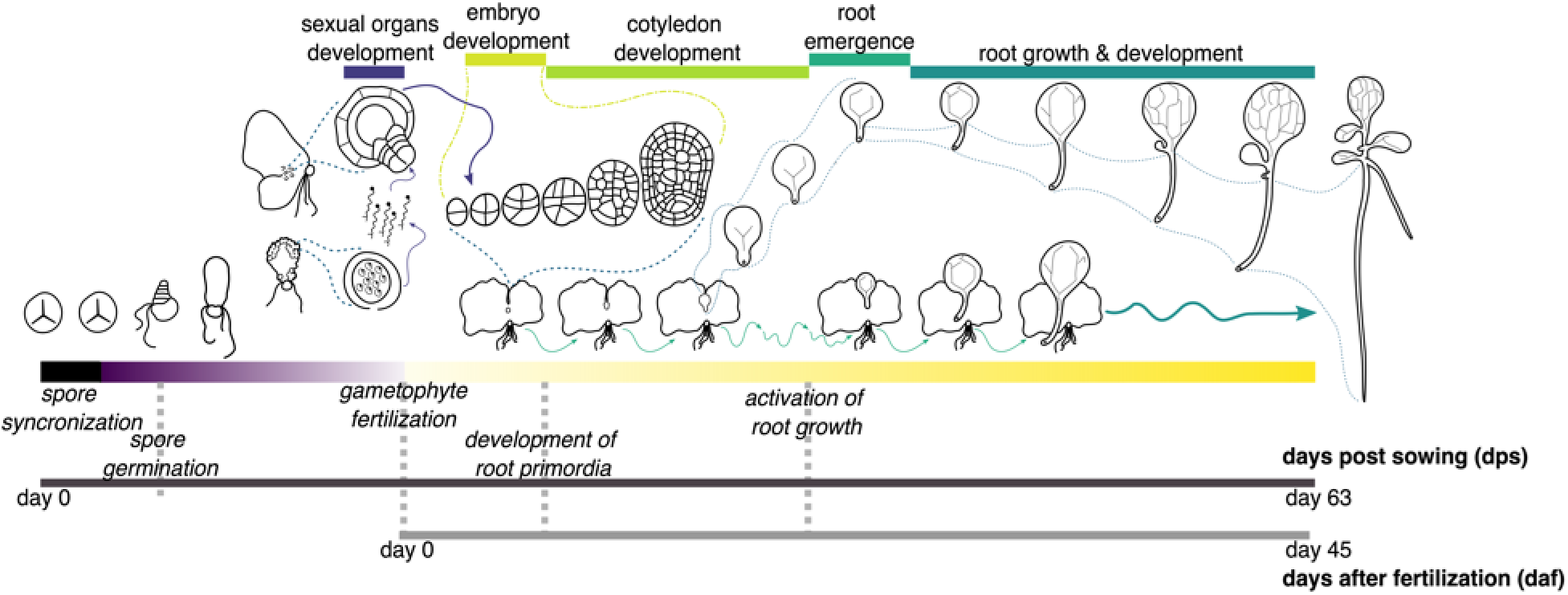
Graphic summary of *Ceratopteris richardii* development from a spore to the young sporophyte, with a focus on the first stem-borne root emergence and development.

### Zonation of the first shoot-borne root

Roots can be divided in three different zones: meristematic (MZ); elongation (EZ); and differentiation zones (DZ). The MZ is located at the distal part of the root where cells are actively dividing with a high mitotic rate. Zonation can be established by measuring the meristem size based on the number of cortical cells from the stem cell niche to the last cortex cell without any signs of elongation (Perilli and Sabatini, 2010; Huang and Schiefelbein, 2015). In the fern *Azolla filiculoides*, this approach was already applied by counting cells in the outer cortex layer (de Vries et al., 2016). We used the same method to determine the different zones of the Ceratopteris root. We detected the outer cortical layer in roots of 30 daf sporelings (**Figs. 2AD**, **4A,B**). Changes in cell length and width are easy to detect in the outer cortex, because this layer does not display any other evident changes during differentiation (**Fig. 4B, dotted line**).

**Figure 4.**
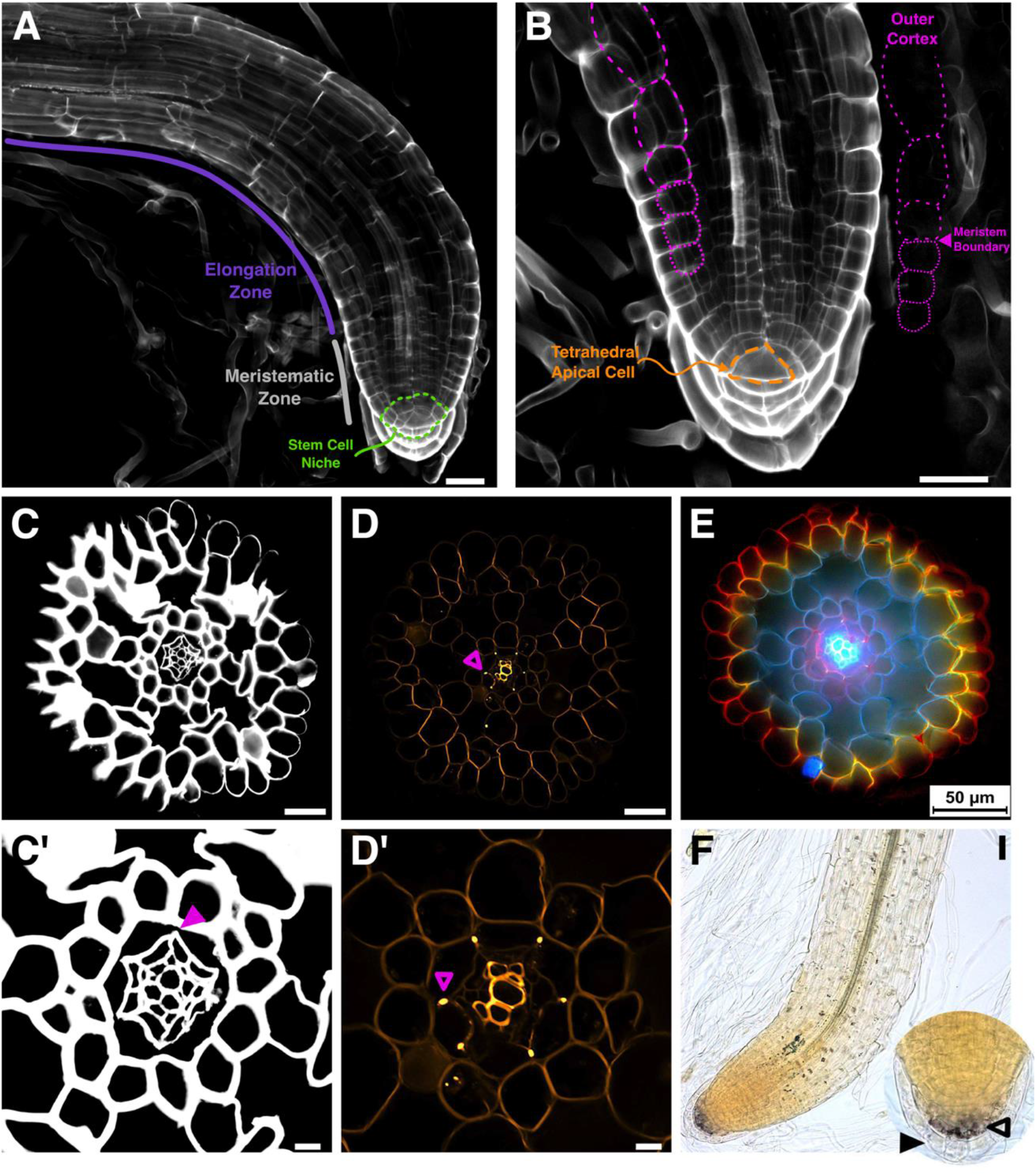
Histological analyses of the *Ceratopteris richardii* first stem-borne root. (**A & B**) Whole root stained with Calcofluor White. The RAM in 30 daf Ceratopteris sporelings has a prominent tetrahedral apical cell. The cell length changes between MZ and EZ after the meristem boundary. This difference is marked with two different dashed lines in purple. (**C & C’**) Transverse section of a root, stained with Calcofluor White to detect cell walls and observed with CLSM. Endodermal cell walls did not stain (purple arrowhead). (**D & D’**) Transverse section of a root stained with berberine and aniline blue. Vasculature and endodermis positions were assigned due to the fluorescent signals in the xylem cell walls and the Casparian strip, respectively (purple unfilled arrowhead). (**E**) Transverse section of a root stained with berberine, lactic acid, and safranin, observed in an epifluorescence microscope. (**F**) Amyloplasts detection using lugol staining in the root. *(Inset)* A strong signal is detected in the middle layer of the root cap (black-filled arrowhead) but not in the outer layer (black-empty arrowhead). *Scale bar: 50 μm (A to F); 10 μm (C’ & D’).*

Cells within the MZ have a similar length and width, while those in the EZ have extended their length. We established the probable meristematic boundary in each root based on discernible cell changes and measured the length of three different cells in each direction, rootward and shootward. We found a distinctive change in cell length among cells inside and outside the MZ boundary (**Table 1A**), where a rapid elongation occurs after cells move away from the meristem. We also assessed cell width, but no considerable change was detected in both zones (**Table 1A**). However, there is a large standard deviation in EZ length due to a rapid elongation change between these cells (**Fig. 4B; wide-dotted line**).

**Table 1.** – Root zonation and meristem size determination of the first stem-borne root of *Ceratopteris richardii*. **(A)** The meristematic zone ends when cells begin to elongate. The change in cell elongation can be detected by measuring length of the outer cortex layer. **(B)** The Ceratopteris first stem-borne root meristem size is seven cells in average.

| A | Cell Size |  |  |  |  | B | Cell Number |  |  |
| --- | --- | --- | --- | --- | --- | --- | --- | --- | --- |
|  | Length (μm) |  | Width (μm) |  | n |  | Mean | SD (+/-) | n |
|  | Mean | SD (+/-) | Mean | SD (+/-) |  |  |  |  |  |
| Meristematic Zone | 16.34 | 2.42 | 17.95 | 2.28 | 20 |  | 7.00 | 1.34 | 50 |
| Elongation/<br>Differentiation Zone | 31.53 | 12.09 | 25.0 | 3.62 |  |  |  |  |  |

After identifying the meristem boundary, based on the number of outer cortex cells, we determined the meristem size, from one-celled merophyte till the last cortical cell at the MZ border. Merophytes were considered because they generate all different proximal tissues, including the outer cortex (**Fig. S3**). We found that the MZ in Ceratopteris first SBR has a median size of 7 cells (SD +/− 1.34, **Table 1B**). Based on these results, the first SBR is composed by a narrow root cap that covers the promeristem or RSCN (**Fig. 4A**), which is inside a short MZ that includes the RAC, merophytes (pluripotent derivatives) and transit-amplifying cells (**Fig. S3**). Ceratopteris meristem size may change because of the root growth dynamics or the effect of diverse stimuli. Our results pretend to establish standardized measures in Ceratopteris first SBR with an active meristem that would be helpful for subsequent analyses in this fern.

### Histochemical identification of the different root cell layers

After longitudinal zones were established, we analyzed the radial arrangement and identity of specific traits within the defined concentric layers of Ceratopteris roots. We obtained transversal sections from the elongation-differentiation root zone (EDZ) of 30 daf sporelings. First, we used calcofluor white (CW) to observe the whole radial morphology of Ceratopteris root, since CW binds to cellulose present in cell walls (**Figs. 4C,C’;** Eeckhout et al., 2013; Huang and Schiefelbein, 2015). We identified six cell layers organized in uninterrupted concentric circles and the central cylinder (**Fig. 4C**). Based on cellular morphology, and a previous report, the central cylinder contains the vascular tissues (xylem and phloem) with the next two layers probably corresponding to the pericycle and the endodermis (**Fig. 4C’**; Hou and Hill, 2004). Subsequently, we performed berberine and aniline blue staining that allows dyeing lignin and suberin that accumulates in endodermis and xylem (**Figs. 4D,D’;** Scheres et al., 1994). We detected the Casparian strip (CS) as highly fluorescent dots between cell walls of the fifth cell layer, showing that this layer corresponds to the endodermis (**Figs. 4D,D’, purple empty arrowhead**). In CW-stained roots, there is no staining of the CS because of the larger accumulation of lignin and suberin, replacing the typical polymers in cell walls which CW is able to bind (**Fig. 4C’; purple filled arrowhead**). We also detected a fluorescent signal in the central cylinder, which may correspond to xylem cells due to their strong lignification and their anatomical characteristics (**Fig. 4D’**).

A previous report showed that the staining of epidermis and exodermis in onion roots after being treated with berberine and safranin (Lux et al., 2005). We choose this procedure to discern the root epidermis and to demonstrate whether an exodermis was present in Ceratopteris roots. We obtained a remarkable staining pattern, since the major cell layers were stained in different colors (**Fig. 4E**). Vascular tissues are detected in a bright blue color, pericycle in pink, endodermis in purple with intense Casparian strip fluorescing dots (**Fig. 4E**). The middle cortex and cortical aerenchyma are dark blue, whereas the outer cortex displays a color gradient, from dark blue (inner cell wall) to yellow (outer cell wall). Finally, the epidermis is stained in bright red (**Fig. 4E**). No sign of an exodermis layer adjacent to the epidermis was found. This result presented a fascinating differential color pattern to distinguish between all root cell layers.

A well-known trait of the differentiated root cap layers (RC), in other species, is the presence of amyloplasts. Therefore, we tested if these types of plastids were present in Ceratopteris RC by using lugol staining (**Fig. 4F**). We observed amyloplasts in the RC cells but also in other cells in the root body at the EDZ. Amyloplasts were observed after the RCI, where root cap cells differentiate (**Fig. 4F; inset, empty black arrowhead**). On the other hand, the last RC layer lacks any stain trace, suggesting that amyloplasts are lost in these cells and/or represents another stage of this layer (**Fig. 4F; inset filled black arrowhead**). Overall, these histological techniques allowed us to distinguish the different cell layers present in the Ceratopteris root: the stem cell niche with the prominent tetrahedral RAC; the amount of lignin in xylem cells depending on their stage; the CS in endodermis; a middle cortex that generates aerenchyma, since Ceratopteris is a semiaquatic plant; the outer radial layers; and different amount of amyloplasts in the protective root cap as a possible marker for differentiation. All these new data can be used in future studies to assess the functional characterization of genes that specify each cell lineage.

### Growth analysis of Ceratopteris stem-borne roots in sporelings

In order to characterize the growth dynamics of Ceratopteris SBRs, we follow root growth on sporelings. Although previous work has been pivotal to understand Ceratopteris root development, we decided to assess growth behavior of the first SBR with respect to the development of further SBRs (Hou and Hill, 2002, 2004). We followed the growth of SBRs after a primordium was noticeable at each node. We recorded root length at diverse time points, defined as days of growth (**Fig. 5**). SBR-1 primordium was present in all sporelings at the beginning of the experiment, but no root elongation was observed until 5 days of growth which could be due to a non-synchronous development (DeMaggio, 1977). At 10 days of growth, SBR-1s of all sporelings showed noticeable growth, SBR-2s were not yet observed. SBR-1 elongation continued slowly for 25 days (**Fig. 5; indigo column**). SBR-1 growth rate (GR) was different from that of the subsequent roots, each newly-emerged root showed faster growth (**Table S2**). SBR-1 displayed a growth rate of 0.17 mm per day from 0-5 days of growth. SBR-2 grew approximately at 0.29 mm/day in a similar 5-days window (10-15 days of growth; **Fig. 5, horizontal lines, Table S2**).

**Figure 5.**
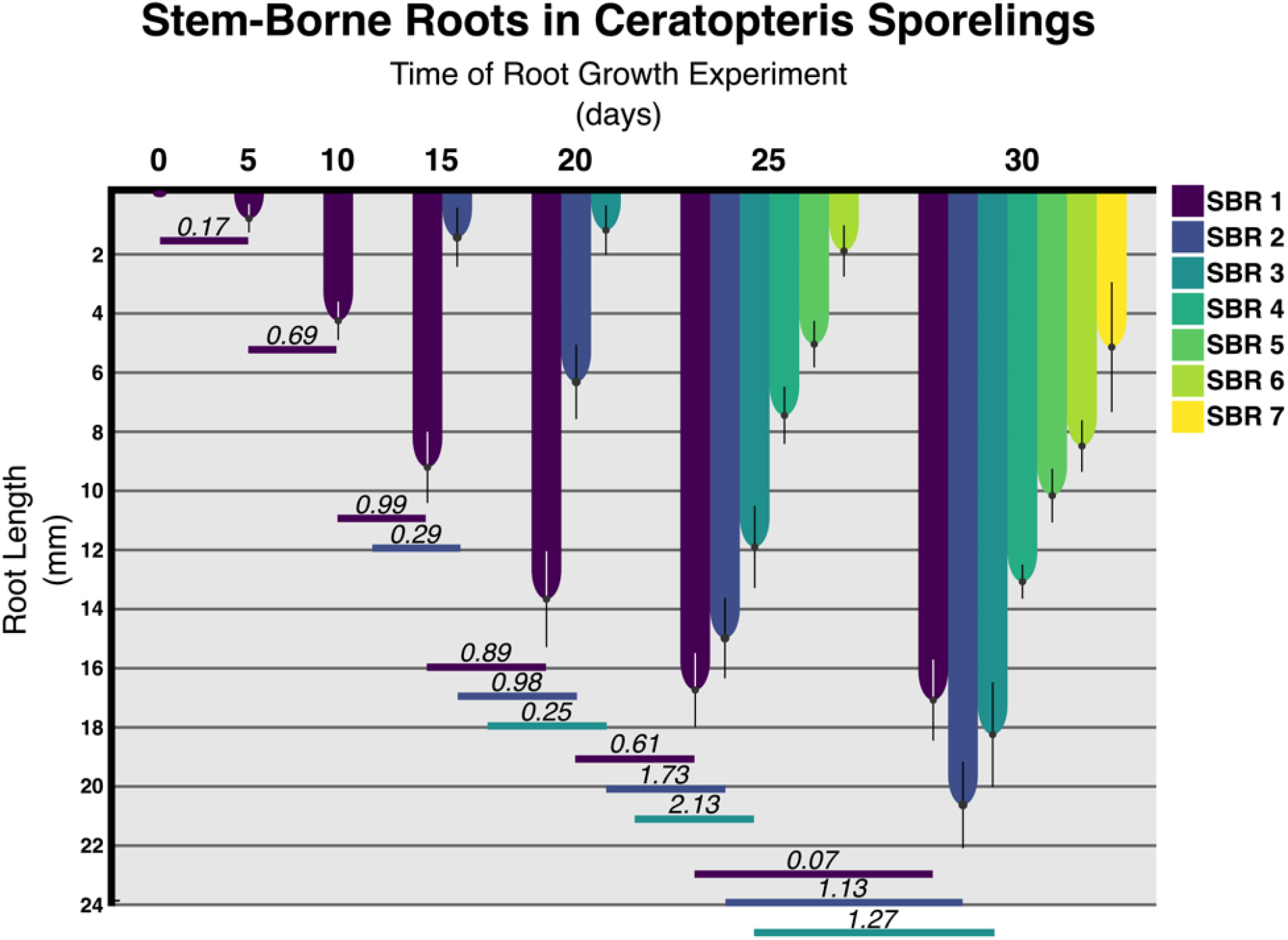
Growth analysis of the stem-borne roots. Column plot showing the growth analysis of stem-borne roots in Ceratopteris sporelings. Horizontal lines describe the growth rate of each SBR (according to color) in a timespan of 5 days. Standard deviation is displayed in each column.

All roots showed a similar behavior in their growth rate pattern through time, initially growing at a slow rate, then increasing the rate continuously until reaching their fastest elongation rate, and finally decreasing their rate. SBR-1 highest GR (0.99 mm/day) was between 10-15 days, after that its GR started to decrease (**Fig. 5; indigo-colored column**). SBR-1 reached its maximum length at 25 days of growth, after which only a marginal growth rate was observed (0.07 mm/day at 30 days **Table S2**). Based on these results, we suggest that SBR-1 enters a determinate growth program after 25 days of growth. The small change in root length at 30 days of growth may not be caused due to an active meristematic growth but to remaining developmental processes, such as cell elongation, as it has been described for other fern roots (Kurth, 1981; Kurth and Gifford, 1985; Nitanyangkura et al., 1980).

The SBR-2 began to elongate in all sporelings from day 15, this root displayed a prominent growth rate from 15 to 25 days of growth (**Fig. 5; tory blue-colored column**). SBR-2 continued an active growth and was longer than the first root at day 30, when its growth rate began to decrease. While SBRs 1 and 2 had a slower emergence time between each other, SBRs 3 to 6 had a prompt emergence in a 5-days timespan (**Fig. 5; days 20 and 25**). Sporelings had the SBR-3 at day 20 and reached an average length of 11.9 mm at day 25 (**Fig. 5; java-colored column**). At day 30, SBRs 4 to 6 were still smaller than older roots, but exhibited active growth and higher growth rates (**Table S2**). On day 25, the seventh root primordium was already present in a few plants, but emerged in all sporelings at day 30 (**Fig. 5; gorse-colored column**).

### Cell cycle activity in the root apical meristem

The RAM of ferns has brought attention over several years, for the presence of the RAC and the precise segmentation patterns that occur in this organ (White and Turner, 1995). Nevertheless, two opposing hypotheses have existed: (1) the RAC acts as a quiescent center and rarely divides; and (2) the RAC frequently enters the cell cycle to divide (Bower, 1889; Gifford, 1983). We determined cell cycle activity in the RAM by exposing Ceratopteris sporelings to 5-ethynyl-2’-deoxyuridine (EdU), after the SBR-1 began an active growth. DNA replication is used as a molecular marker to detect cell cycle progression, and arabinosyl nucleosides allows the detection of cells entering to the S-phase of the cell cycle (Cruz-Ramirez et al., 2013; Fujinami et al., 2017; Neef and Luedtke, 2012). Sporelings were collected at different times of EdU exposure to evaluate cell division activity. We divided Ceratopteris RSCN based on distinctive cell fates and/or developmental stages that could influence cell cycle activity (**Fig. 6G**): the RAC (**indigo-colored cell**); merophytes or proximal initials (**dark green-colored cells**), direct daughters of the RAC involved in formative divisions; proximal cells (**light green-colored cells**), derived directly from merophytes; root cap initial or RCI (**blue-colored cells**), distal derivative of the RAC; and root cap cells (**yellow-colored cells**), distal cells entering a differentiation process.

**Figure 6.**
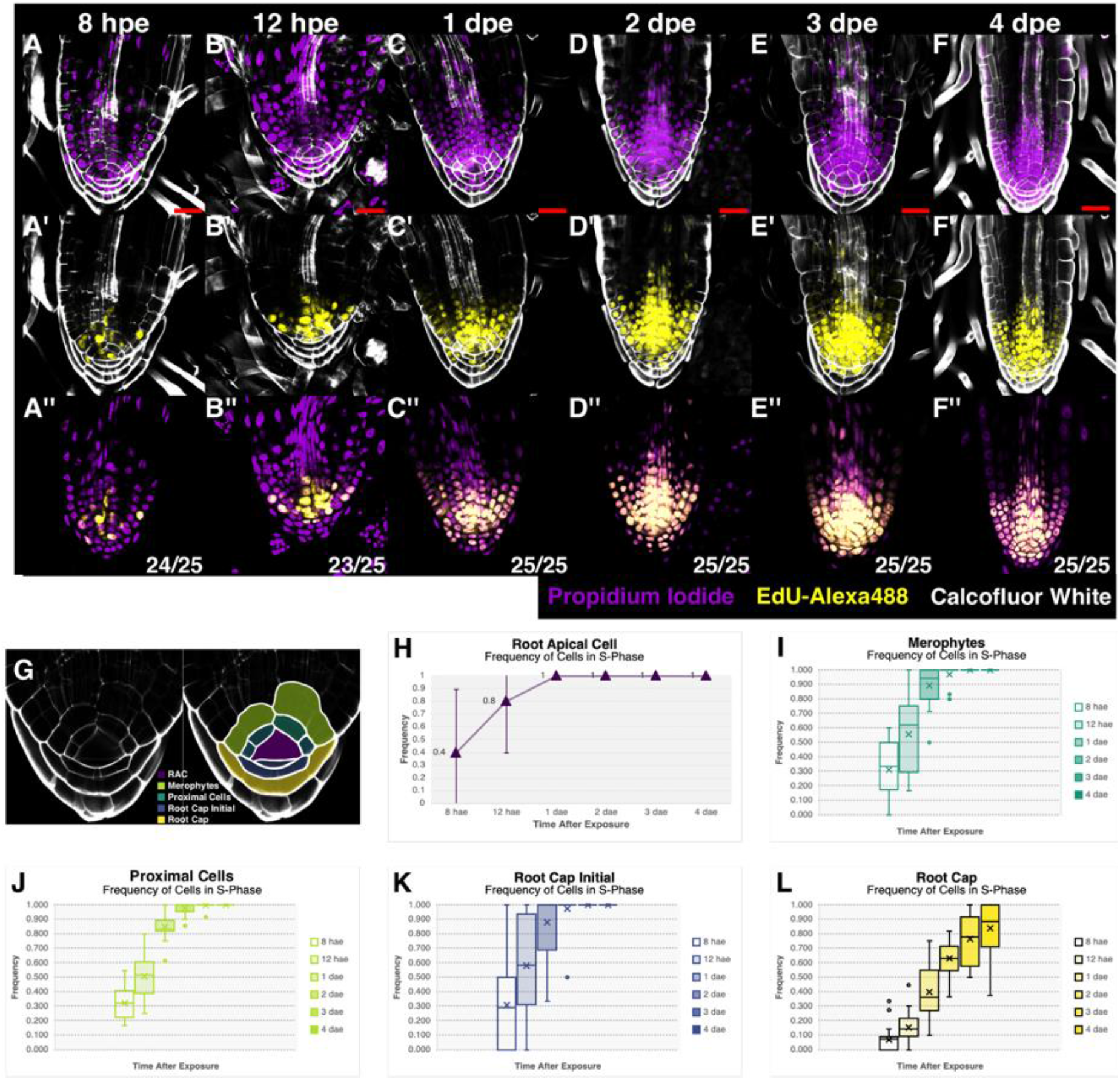
Cellular dynamics of the first shoot-borne root. (**A to F**) EdU incorporation experiment to detect cells in their S-phase of cell cycle at 8 hae (**A, A’ & A’’**), 12 hae (**B, B’ & B’’**), 1 dae (**C, C’ & C’’**), 2 dae (**D, D’ & D’’**), 3 dae (**E, E’ & E’’**), and 4 dae (**F, F’ & F’’**). *(A to F) DNA counterstaining and cell wall staining (Propidium Iodide – Purple; Calcofluor White – White); (A’ to F’) EdU-labeled DNA and cell wall staining (EdU-Alexa488 – Yellow; Calcofluor White – White); (A’’ to F’’) EdU-labeled DNA and DNA counterstaining merge (EdU-Alexa488 – Yellow; Propidium Iodide – Purple). Scale bar: 20 μm. hae – hours after exposure; dae – days after exposure.* (**G**) Established zonation inside the stem cell niche. (**H**) Scattered plot of the frequency of S-phase cells in the RAC according to the exposure time. (**I to L**) Box plots showing the frequency of entering the S-phase of cell cycle in the different zones of the RAM: merophytes (**I**); proximal cells (**J**); root cap initials (**K**); root cap cells (**L**).

Roots exposed to EdU showed a clear incorporation signal in the nuclei of the different cell types analyzed (**Figs. 6A’-F’**). These observations were compared with nuclei counterstaining with propidium iodide (**Figs. 6A-F**). The merged signal between EdU-Alexa488 and PI shows the proper signal position in each nucleus (**Figs. 6A’’-F’’**). Notably, right after 8 hours of exposure, EdU incorporation was observed in 40% of analyzed RACs (**Fig. 6F**) as revealed by their stained nuclei (**Fig. 6A’**). In subsequent exposure times, more roots displayed a stained RAC nucleus (**Figs. 6B’,H**). From 1 dae, all roots analyzed showed incorporation signal in the RAC (**Figs. 6C’-F’,H**). These results suggest that the RAC undergoes constant cell divisions in the SBR-1 of Ceratopteris sporelings. Such developmental behavior of the RAC may indicate that this cell is not mitotically quiescent. However, because of its position in the RSCN, we propose that the RAC acts as an organizing center that receives and produces different signals to coordinate proper organ formation.

Also, merophytes and proximal daughter cells displayed incorporation signals at early exposure times (**Fig. 6A’**). Both types of cells showed a similar incorporation frequency at 8 hae (**Figs. 6I,J**). But at 12 hae and 1 dae, merophytes displayed an incorporation frequency of 10% and 5% faster compared to proximal daughter cells. Merophytes act as progenitors of proximal cells, and they are subjected to repeated rounds of cell division, which may be reflected by a high incorporation frequency (Kurth, 1981; Hou and Hill, 2004). Instead, proximal cells can have a limited number of proliferative divisions depending on the cell layer (**Fig. S3**), which could explain a lower EdU incorporation, compared to merophytes.

The RCI presents a peculiar EdU incorporation pattern (**Fig. 6K**). We detected that the RCI showed more than a nucleus or a continuous cell wall (**Figs. 6A-F**). This may indicate the RCI is subject to proliferative divisions, because we observed up to four different nuclei at this position. The root cap cells exhibited incorporation values from 2 to 4 times slower compared to the other zones in early exposure times (**Fig. 6L**). Also, it was the only cell lineage that did not incorporate EdU in all analyzed cells, which suggests that these cells lose their division capacity while distancing from the RAC, probably reaching a differentiated status.

## DISCUSSION

### How early is the first root specified during Ceratopteris embryogenesis?

By delving into early sporophyte development in *Ceratopteris richardii*, we determined the specification of the first stem-borne root and growth dynamics of its homorhizic root system (**Fig. 3**). The cellular ontogeny of the first SBR has been debated several times. Separate authors proposed different explanations at what stage and from which cell is established (Chiang and Chiang, 1962; DeMaggio, 1977; Johnson and Renzaglia, 2008; Wardlaw, 1955). Our rigorous observations of the different stages in Ceratopteris embryogenesis suggest that the first root is specified, at least, at the octant stage from the hypobasal outer cells (**Fig. 2G**). From those cells, the distinctive tetrahedral-shaped RAC would be distinguishable in later stages (**Figs. 2H,I**) and the whole RAM would develop (**Figs. 2P’, 4A**). These results show that the early patterning *Ceratopteris richardii* embryo is similar to other leptosporangiate ferns (Johnson and Renzaglia, 2009). In Arabidopsis, root development begins by recruiting the most upper suspensor cell, the hypophysis, which would later give rise to the quiescent center and columella. Even though, Arabidopsis RAM is observable until the late globular stage, the hypophysis is already present since the 8-celled embryo based on the overlapping expression of two homeodomain genes from the WOX family, *AthWOX8* and *AthWOX9* (Jenik et al., 2007; Bennett and Scheres, 2010). In Ceratopteris, five different WOX genes were identified, with *CriWOXA* and *CriWOXB* belonging in the same clade as those previously mentioned (Nardmann and Werr, 2012; Liu and Xiu, 2018). No studies assessing WOX genes in Ceratopteris embryogenesis have been performed yet (Nardmann and Werr, 2012; Youngstrom et al., 2019; Yu et al., 2020). Transcriptional analyses, transgenesis and reverse genetics approaches in Ceratopteris are necessary to detect key regulators during the embryogenesis, to uncover how early the specification of the root is carried out, and if regulatory networks in this process are conserved between seed plants and ferns.

### The Ceratopteris root develops from a narrow multicellular meristem

The discrete root zonation exhibits the developmental and cellular processes carried out in each sector. This feature has been well documented from Arabidopsis and other angiosperms to pteridophytes (de Vries et al., 2016; Huang et al., 2015; Perilli and Sabatini, 2010). We established the different root zones in the first SBR of Ceratopteris (**Figs. 4A, 7A**), which is composed of a small-sized meristematic zone (**Table 1B**) followed by the EDZ. A straightforward separation between the EZ and DZ was not determined: root hair development was not observed right after the MZ but xylem differentiation was noticed (**Fig. 4A, purple line**). This ambiguous partition between both zones could be a consequence of the developmental stage when Ceratopteris roots were analyzed. According to analysis in other fern species, *Azolla filiculoides* roots displayed the three different zones, but these roots were produced from a regenerative process (de Vries et al., 2016). Additionally, Azolla roots presented a larger meristem size (**Fig. 7B, middle panel**) when compared to Ceratopteris roots **Fig. 7B, left panel**).

**Figure 7.**
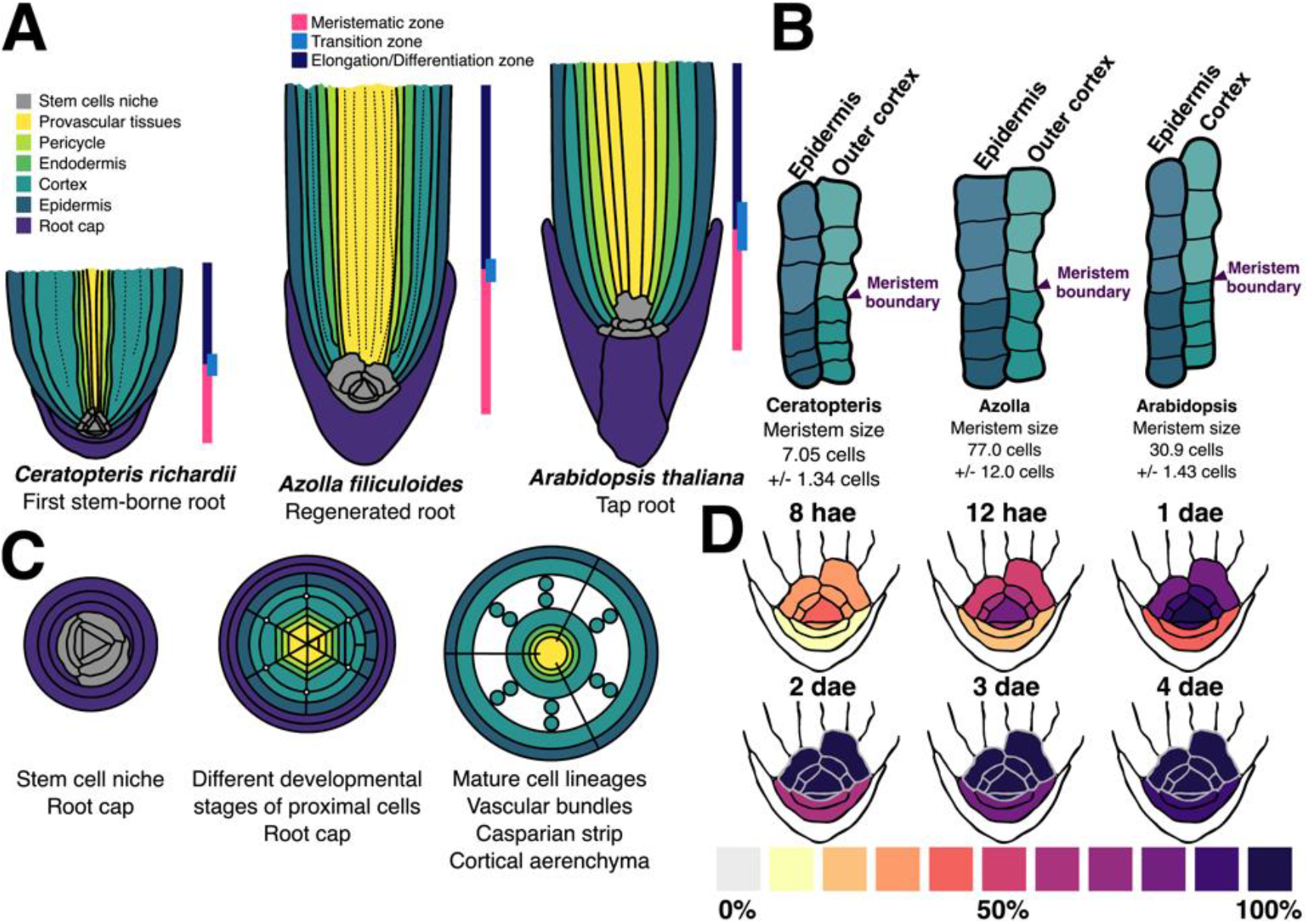
Graphic summary of the cellular organization and division dynamics in the *Ceratopteris richardii* first stem-borne root. (**A**) Roots are divided in three zones: meristematic, elongation, and differentiation zone. Ceratopteris root presents two main zones: meristematic and elongation/differentiation zones. The roots of Azolla and Arabidopsis are represented for comparison purposes. (**B**) In a longitudinal section, the boundary between meristematic and elongation/differentiation zones is established where the cell length begins to expand. For Azolla and Ceratopteris, the outer cortex layer has a reliable morphology. The size of root apical meristem varies considerably among the three species depicted here. (**C**) Radial organization changes according to the developmental stage. (Left) The RAC and its merophytes are surrounded by root cap cells. (Middle) Cell transitions from merophytes to specific layer initials. (Right) Differentiated tissues with specific traits. **(D)** Different cell types of the RAM have different cell cycle activity, based on EdU incorporation analysis. The RAC has a higher activity compared to its derivatives.

Furthermore, Arabidopsis seedlings display a root meristem with an average size of 30.9 cells (**Fig. 7B, right panel**; Perilli and Sabatini, 2010) which could be considered an intermediary from the meristem sizes in fern roots. But Arabidopsis is a phylogenetically distant plant and even its meristem organization is divergent from ferns and other angiosperms. The dramatic difference in size of the Ceratopteris MZ, 11 and 4 times smaller than Azolla’s and Arabidopsis’ MZ, respectively, opens up the question of how the meristem size is regulated. The phytohormones, auxin and cytokinin, have shown opposing roles in controlling the meristem size in Arabidopsis (Perilli and Sabatini, 2010). Also, the activity of the proteins PHABULOSA (PHB) in the stele and SCARECROW (SCR) in the endodermis have been linked to modulate meristem size by controlling the cytokinin response regulators in the proximal meristem, thus affecting the number of dividing cells and root growth (Moubayadin et al., 2016; Sebastian et al., 2015). Recent studies reported an orthologous gene for *SCR* and the presence of *PHB* gene subfamily (Class III HD-Zip, Type 2) in several fern species (Motte et al., 2019; Vasco et al., 2016). Still, the response to hormones in ferns roots doesn’t seem to be similar and even opposed to Arabidopsis, which places a challenge regarding the role of these genes in controlling the root meristem size (de Vries et al., 2016; Hou et al., 2004; Yu et al., 2020).

The Ceratopteris RAM is composed by the RAC, the proximal merophytes and a distal RCI. This arrangement is maintained since the RAC appearance during embryogenesis to postembryonic development (**Figs. 2M,4A,7A**). Our morphological analyses showed a consistent organization compared to previous reports on Ceratopteris (Hou and Hill, 2002; 2004). Also, this meristem organization is highly conserved in roots of different fern species (**Fig. 1B, Table S1**). Based on our observations, and those reported for other ferns, we suggest that the RAC and its derivatives comprise the stem cell niche of the Ceratopteris root. Still, more experiments are needed to determine hormone gradients, expressed genes and other cues implicated in RSCN activity and maintenance. Noteworthy, exceptions of this consensus morphology are present in other fern species. In some eusporangiate ferns and Osmundales, the RAM has two to four (smaller) apical cells instead of a single RAC (Freeberg & Gifford, 1984; Gifford, 1983; White and Turner, 1995).

Our proposed fern RAM organization contrasts with the RAMs from other major lineages of vascular plants, which exhibit higher morphological diversity. For example, in lycophytes four different types of RAM have been defined, depending on their cellular organization and division frequency (Fujinami et al., 2017). Similar to our findings, the RAMs in the order Selaginellales also have an apical cell at their tip (Otreba and Gola, 2016). The presence of this RAM organization in Selaginellales and several orders of ferns could imply different scenarios: (1) meristems with an apical cell were present in the ancestor of vascular plants, later co-opted for root meristems and diversified into a wide range of cellular organizations except for ferns and Selaginellales (**Fig. 1B**); (2) this RAM organization appeared separately in ferns and Selaginella species, and it has been preserved because of its efficient physiology. How an increase in diversity can be gained in the RAMs at the cellular level still remains unanswered and becomes a fascinating subject to unravel specific adaptations at a cellular level.

### Ceratopteris root cell layers exhibit the conservation of specific traits

A root is composed by a diverse set of cell layers, radially organized, each one with its own identity and function (Bennett and Scheres, 2010; Dolan et al., 1993). We focused mainly on detecting specific traits that appear during the maturation of each cell layer and used Hou and Hill (2004) report as a reference to examine our results. In this aforementioned study, the transversal organization of the RSCN was uncovered with the RAC and merophytes (from different cell numbers) were at the center and surrounded by root cap cells (**Fig. 7C, left**). This organization is localized at the most distal part of the MZ, where merophytes have only been subjected to a couple of proliferative divisions (Gunning et al, 1978; Hou and Hill, 2004). A crucial developmental stage comes after the former (shootwards), and there, the merophytes continued dividing (**Fig. 7C, middle;** Hou and Hill, 2004). The proximal merophytes can be considered transit-amplifying cells as they are still inside the MZ and dividing several times in a precise pattern to generate the diverse cell lineages of the root body (**Fig. S3**). Upon entering the EDZ, we detected a clear concentric organization of the cell layer and each one of these displayed specific traits, which allows the distinction from one another (**Fig. 7C, right**). The diverse set of features analyzed in the EDZ of Ceratopteris roots seems to be conserved with other vascular plants (Barlow, 2003; Bennet and Scheres, 2010; Geldner, 2013; Heimsch and Seago, 2008; Imaichi et al., 2018; Jung et al., 2008; Kirschner et al., 2017; Lucas et al., 2013; Pires and Dolan, 2012; Takahashi et al., 2014). This could imply that some cell lineages already existed in the common ancestor of euphyllophytes, and suggests the conservation of genetic networks involved in their specification, to some extent, between ferns and seed plants.

The vascular tissues were an important innovation of tracheophytes, they are divided in two different functional domains: xylem and phloem (Pittermann et al., 2015). We detected the presence of lignin at the center of Ceratopteris roots, which demonstrate the localization of the xylem. Additionally, a differential signal intensity and cellular morphology made discernable xylem types: metaxylem, higher staining signal and wider cells; and protoxylem, minor signal and smaller cells. This could be a consequence of a greater lignin accumulation due to an advanced maduration state (De Rybel et al., 2015; Lucas et al., 2013). We did not detect a specific trait of phloem, but unstained cells adjacent to xylem cells correspond to this tissue. Another study reported two different antibodies binding preferentially to pectin or hemicellulose present in the phloem, and that can be used in future research to differentiate this layer (Eeckhout et al., 2014).

The endodermis bears a Casparian strip, which is composed by lignin and lower amounts of suberin (Geldner, 2013). We observed the CS at the center of transversal and anticlinal cell walls in the fifth cell layer (**Fig. 4D’**). The CS was previously observed in other Ceratopteris species and different ferns (Chiang and Chou, 1974; PriestleyI and Radcliffe, 1924). The endodermis has been constantly present in euphyllophytes roots for millions of years, probably due to its function in water and nutrient transport. Interestingly, the roots of Lycopodium species (Lycodiopsida) were reported to lack a CS, thus neither an endodermis (Damus et al., 1997).

The root cap is a common feature of vascular plants. This tissue is considered a main innovation in roots because it protects the RSCN (Kumpf and Nowack, 2015). We detected the presence of amyloplasts in the root cap zone of Ceratopteris, and low signal was also observed within the root apex (**Figure 3F**). Amyloplasts have been found in other ferns, in the root cap of *Regnellidium diphyllum* and in the stele of *A. filiculoides* roots (de Vries et al., 2016; Eastman and Peterson, 1985). Noteworthy, a recent report concluded that the role of amyloplasts in gravitropic response is absent in the roots of ferns and lycophytes (Zhang et al., 2019).

While some studies have begun to unravel the developmental role of certain genes in Ceratopteris and other ferns, none of them focused on root cell layers specification (Ambrose and Vasco, 2016; Bui et al., 2017; Cruz et al., 2020; Hasebe et al., 1998; Plackett et al., 2018; Sano et al., 2005; Vasco et al., 2016; Youngstrom et al., 2019; Yu et al., 2020; Zhang et al., 2017). Unsurprisingly, the genetic networks involved in root cell layers have been mainly described in Arabidopsis (Bennett and Scheres, 2010; Kumpf and Nowack, 2015; Motte et al., 2020). We already speculated the possible or partial conservation of gene function in cell layer specification of ferns and angiosperms roots. One example in Ceratopteris root might be the specification of middle cortex and endodermis from a common merophyte derivative (**Fig. S3**). In Arabidopsis root, a single initial is present for both ground tissue layers, where *AthSCR* and its partner *SHORTROOT* (*AthSHR*) have been widely characterized as essential effectors in the specification process (Bennett and Scheres, 2010; Motte et al., 2020). Their orthologous genes are expressed in Azolla root tip, which could indicate the conservation of this network, considering more genes are involved (de Vries et al., 2016). Fern orthologs for other genes expressed during Arabidopsis root development have been reported: the *Class III HD-Zips* and *VND* genes involved in xylem differentiation were detected in Azolla root and the leaves of other fern species (de Vries et al., 2016; Vasco et al., 2016); *KANADI* genes implicated in phloem development were identify in *Equisetum hyemale* leaves (Zumajo-Cardona et al., 2019); the NAC domain genes, *FEZ* and *SOMBRERO*, required for root cap development, were detected to be preferentially expressed in the MZ of six different plants, including the lycophyte Selaginella (Huang et al., 2015). Still, not all structural orthologs found in Ceratopteris have the “complete” function from their equivalents in Arabidopsis, such as *CriWUL* and *CriPINJ*, a *WUSCHEL* ortholog and the sole ortholog for the extensively duplicated Eu3-PIN clade (Bennett et al., 2014; Zhang et al., 2017; Zhang et al., 2019). The specification of cell layers poses the possibility to explore the conservation and divergence of the genetic networks that define cell lineages.

### Ceratopteris stem-borne roots display growth cessation

Ceratopteris establishes a complex organization of its homorhizic root system throughout development. Hou and Hill (2002) followed the growth behavior of six SBRs and focused on determining their heteroblastic development and growth rate. Our approach explored the emergence and growth of Ceratopteris SBRs for 30 days after the first root emerged (**Fig. 5**), while examining the timeline of the root system establishment and growth behavior in solid media. We detected that root emergence time is shortened between consecutive SBRs. We propose two possible explanation for this phenomenon: (1) a wider leaf area available for photosynthesis would produce higher amounts of photosynthates available to develop other organs; (2) a more complex root system would allow a more efficient water and nutrient uptake, allowing faster organ formation. We observed the cessation of root growth, following a decrease in growth rate, in the earliest SBRs compared to recently-emerged roots. Hou and Hill (2002) found a similar outcome while Ceratopteris plants growing in liquid media, which lead us to conclude that media firmness has little to none effect on root growth.

The termination of root growth suggests that Ceratopteris SBRs follow a determinate growth program, even in the presence of nutritional sufficiency. This is concordant with the concept of primary homorhizy indicating that, at least, the first SBR has a short time span (Schneider, 2013). Determinate growth has been shown in different groups of angiosperms. Some cacti species display a constitutive determinate program in their primary root, that is considered an adaptation to severe drought. This process is characterized by the spontaneous RAM exhaustion: reduction of the proximal meristem; root hair development at the root tip; increased lateral root formation; terminal differentiation of the RSCN. The *Stenocereus gummosus* roots (Cactaceae) have been studied to understand the molecular mechanisms underlying a determinate program. This developmental process can be induced in Arabidopsis roots by nutrient deficiency, redox state and other detrimental stimuli. The root meristem exhaustion in *Stenocereus* recapitulates the developmental zonation of Arabidopsis. *Stenocereus* roots at terminal stage displayed a similar transcriptional profile to the differentiation zone of Arabidopsis, where genes involved stemness and proliferation are downregulated (Benesova, 2016; Gutierrez-Alanis et al., 2015; Rodriguez-Alonso et al., 2018; Rodriguez-Rodriguez et al., 2003).

Azolla roots also showed a determinate program, the RAC was found to divide around 55 times. No morphological changes were reported in mature Azolla roots besides a progressive diminution of plasmodesmata connections (Gunning, 1978). The RAC is mentioned as a consistent trait of the RAM in young and mature roots (Gunning et al., 1978), which supports our hypothesis of the RAC as an organizing center. The omnipresence of the RAC in mature roots of *Equisetum scirpoides* and *Marsilea vestita* have been indirectly reported (Gifford and Kurth, 1982; Kurth, 1981). This could suggest that the root determinate program does not display major morphological changes and is a consensus behavior in ferns, including Ceratopteris. Still, further research should evaluate what leads the RAC towards an inactive cell cycle, the mechanical importance of the RAC shape even in the absence of new divisions, and the overall developmental and physiological changes that define root determinacy in Ceratopteris.

### The Ceratopteris RAC exhibits high mitotic activity

The RAM is considered a self-preserving tissue that defines the continuous development of different tissues (Fujinami et al., 2020). Our analysis of Ceratopteris root meristem dynamics demonstrated that the RAC has a high division frequency because of signal detection at early treatment times (**Fig. 7D [8 hae]**). This indicates an active cell cycle in the RAC and a higher activity than its immediate derivatives (**Fig. 7D, upper panel**), which is consistent with the RAC histogenetic role (Hou and Hill, 2004). Nevertheless, other studies have concluded that the RAC in ferns is mitotically active and progress throughout the whole cell cycle by looking mitotic figures and using colchicine to avoid mitosis progression (Chiang, 1972; Gunning et al., 1978; Nitayangkura et al., 1980). Merophytes are essential cellular units for the root proper formation and the immediate derivatives of the RAC. (Barlow, 1989). The proximal merophytes and the RCI showed a slower incorporation frequency compared to the RAC at the earlier exposure times (**Fig. 7D, upper panel**). A similar observation was reported in Marsilea and Azolla, where the RAC displays a higher cell division frequency compared to the merophytes (Kurth, 1981; Kurth and Gifford, 1985).

The dynamics observed in Ceratopteris RAM offers the possibility to assess how genetic networks and different signals operate to maintain a constantly dividing stem cell niche. Auxins were found to play an important role during root initiation in Ceratopteris. Genes involved in the cell cycle, DNA replication and chromatin remodeling are upregulated after one day of treating Ceratopteris sporelings with exogenous auxins. Orthologous genes for the WOX, GRAS and PIN gene families, well-known RSCN regulators, were also activated (Bennett and Scheres, 2010; Yu et al., 2020). *CriWOXA* was found as a direct target from auxin response and transiently expressed in the root apical mother cell. CriWOXA will later active *CriWUL* expression in one-celled merophytes (Nardmann and Werr, 2012; Yu et al., 2020). But both genes were not expressed in the same cell. A possibility is CriWOXA movement to adjacent cells to form a protein gradient that allows the activation of *CriWUL* and other genes. An Arabidopsis ortholog behaves in this manner: *AthWOX5* is expressed in the QC and its protein moves towards stele and columella initials (Yu et al., 2017). Still the presence of hormone gradient and the actual roles of genes involved in Ceratopteris root development and RAM dynamics remain to be elucidated.

A fine-tune explanation for each developmental process observed in our results demands an extended knowledge of *Ceratopteris richardii*. Whilst we provided a comprehensive description of Ceratopteris stem-borne roots during their development, collectively with previous studies (Conway and Di Stilio, 2020; Hou and Hill, 2002), we have contributed to a more complete framework of Ceratopteris developmental stages. Hence, this fern could be used to assess the cellular and molecular mechanisms in fern root development that remains unexplained. Also, Ceratopteris stands as a major model plant for evo-devo studies due to its evolutionary position in the plant phylogeny (Delaux et al, 2019). Thus, ferns have entered the modern era of plant science, with Ceratopteris as a compelling sidekick to explore the diverse evolutionary paths that this lineage have undergone in their history on Earth.

## Material and Methods

### Plant growth and culture conditions

Ceratopteris Hn-n spores were cultured in C-Fern Medium (CFM, 0.5 g/L MES, pH 6.0, 0.8% agar) at 25°C with a photoperiod of 16h/8h light/dark. Spores were sterilized prior to culture (Hickock et al., 1995) and synchronized in darkness for three days. When gametophytes were found sexually competent (15 days post sowing [dps]), water was added to the culture for the fertilization to take place. In the following experiments, 45 dps sporelings (visible cotyledon and root primordium) were cultured in CFM plus 2% sucrose at the same conditions.

While previous works stated 28 °C as the optimal growth temperature, Ceratopteris can also grow from 21 to 30 °C. Ceratopteris development would increase up to twice at 21-22 °C compared to optimal temperature (Hickok et al., 1995; Hickok and Warne, 1998; Warne and Lloyd, 1980). An extended duration of Ceratopteris development was observed in our results because of the growing temperature used in this work (25 °C). But no morphological change was observed when compared with previous reports.

### Fixation and clearing

Gametophytes and sporophytes were fixed immediately after collection in 4% p-formaldehyde and 0.1% Triton X-100 in PBS 1X for 30 minutes and vacuum was applied. Following, tissues were washed using PBS 1X. All tissues were cleared in the ClearSee solution for at least one week. ClearSee was changed in sporophyte tissues every three days. We used the ClearSee technique since it allows the visualization of the whole plant (Kurihara et al., 2015).

### Tissue sections and staining

Plant sporelings were embedded in 4% agarose and hand-sectioned using razor blades. For transversal staining, 1ug/mL Calcofluor White (or Fluorescent Brightener 28 [Sigma – F3543] was applied to agarose sections for one-Four prior the observation (Huang and Schiefelbein, 2015). For Casparian strip detection, fresh root sections were treated as Scheres et al., (1995). For differential layer detection, fresh root sections were treated with a modified version as Lux et al., 2005; agarose sections were incubated in lactic acid for one hour, then transferred to berberine hemisulfate solution (0.1% in lactic acid) and incubated for 90 minutes at room temperature. Sections were washed with water until they were cleared and then incubated in 0.5% safranin O (in 50% ethanol) less than a minute. Finally, sections were washed with water until cleared. All slides were mounted in 50% glycerol.

For observation of amyloplasts, whole roots were cleared in ClearSee (Kurihara et al., 2015) and then stained in lugol solution. Also, Ceratopteris sporelings were stained with 1 ug/mL Calcofluor White (CW) and 20 ug/mL Propidium Iodide (PI), to observe cell wall and nuclei, respectively. Both dyes were applied in ClearSee. First, CW for 1 hour, then wash in ClearSee for 30 minutes. Second, PI for 30 minutes and then wash in ClearSee for 30 minutes. Finally, samples were mounted in 50% glycerol. This treatment was also applied for gametophytes.

### Root growth analysis

Sporelings of 45-dps were transferred to CFM plus 2% sucrose and 0.4% Gellan Gum (G434 – PhytoTech Labs) and placed vertically in a growth chamber at the same aforementioned conditions. Two media plate changes were applied to each original plate (once each fifteen days). Each sporeling was handled carefully during change of media plate, assuring that roots were in contact with the solid medium. To avoid undesired effects due to contamination, nutrient limitations, agar dehydration and space constraints, we carefully changed sporelings to fresh growth medium every ten days. Root growth was registered every five days in each visible root or organ primordia observed for 30 days. We monitored the development of the earliest seven SBRs (SBR1-7), until extra root primordia appeared at the base of the seventh leaf (leaf-borne roots). The sample was composed of 30 sporelings and for subsequent analysis we randomly selected 20 of these sporelings.

### EdU incorporation treatment

For the EdU incorporation, 45-dps sporelings were cultivated in CFM plus 2% sucrose and 4 mM EdU plates. Sporelings were collected and fixed as previously mentioned at different times (8 and 12 hours, 1, 2, 3, 4 and 5 days) after culture. Sporelings were cleared in ClearSee for one week, stained in CW for 1 hour, washed in PBS 1X two times for 10 minutes, then the Click-iT EdU staining kit (C10337, Invitrogen) was used for signal development, washed in PBS 1X two times for 10 minutes and stained in PI for 30 minutes, washed in ClearSee for 30 minutes and mounted in 50% glycerol, prior their observation (Cruz-Ramírez et al., 2013; Neef and Luedtke, 2011).

### Microscopy and image analysis

Confocal laser scanning microscopy (CLSM) images of both gametophytes, sporelings, and root sections were performed on a Zeiss LSM800 microscope. The different dyes used were excited and detected as follows: calcofluor white, ex: 405 nm laser, dt: SP 470 filter band; Alexa488, ex. 488 nm laser, dt. SP 545 filter band; propidium iodide, ex. 561 nm laser, dt. LP 575 filter band; berberine hemisulfate and aniline blue, ex. 488 nm laser, dt. SP 545 filter band. All lasers were used at 15% intensity and detected using 1 pinhole unit. Differential interface contrast microscopy was used to take whole-mount roots stained with lugol and other root sections, on a Leica DM6000B microscope. Using this microscope, sections stained with berberine hemisulfate and safranin were subjected to fluorescence light and images detected while using a GFP filter. The gametophyte-to-sporophyte development images were taken on a stereo microscope Leica EZ4-FD. Images were organized using Inkscape 0.92.4. Line art figures were designed using Inkscape 0.92.4 and colored with GIMP 2.10.12.

## Abbreviations

RAM: Root Apical Meristem
RAC: Root Apical Cell
RCI: Root Cap Initial
RSCN: Root Stem Cell Niche
SBR: Stem-Borne Root
QC: Quiescent Center
dps: days post sowing
daf: days after fertilization
hae: hours after exposure
dae: days after exposure

## Acknowledgments

We thank Jane Langdale and Andrew Plackett (Oxford University), and Pamela Soltis, Doug Soltis and Blaine Marchant (University of Florida), for providing us with *Ceratopteris richardii* Hn-n spores. To our lab manager Annie Espinal-Centeno for obtaining the first batch of Ceratopteris spores to begin with this project.

## Authors Contributions

A.A-R., L.H-E. and A.C-R. conceived the project and generated the experiment design. A.A-R. performed all experiments. A.A-R., I.B., L.H-E. and A.C-R. analyzed the data. All authors helped in the interpretation of results. A.A-R. and A.C-R. wrote the manuscript with inputs from all co-authors.

